# Welfare at group and individual level: optical flow patterns of broiler chicken flocks are correlated with the behaviour of individual birds

**DOI:** 10.1101/2021.01.19.427267

**Authors:** Sabine Gebhardt-Henrich, Ariane Stratmann, Marian Stamp Dawkins

## Abstract

Group level measures of welfare such as the optical flow patterns made by broiler chicken flocks have been criticized on the grounds that they give only average measures and overlook the welfare of individual animals. However, we here show that by using the skew and kurtosis in addition to the mean, optical flow patterns can be used to deliver information not just about the flock average but also about the proportion of individuals in different movement categories. We correlated flock optical flow patterns with the behaviour of a sample of 16 birds per flock in two runway tests and a water (latency-to-lie) test. In the runway tests, there was a positive correlation between the time taken to complete the runway and the skew and kurtosis of optical flow on day 28 of flock life (slow individuals came from flocks with a high skew and kurtosis). In the water test, there was a positive correlation between the length of time the birds remained standing and the mean and variance of flock optical flow (the most mobile individuals came from flocks with the highest mean). Patterns at flock level thus contain valuable information about the welfare of the individuals that compose the flock.

**Simple Summary:** Technology on farms potentially brings benefits of improved animal health, welfare and productivity as well as reduction in disease, waste and environmental impact. However, it also raises public concern about the welfare of individual animals, particularly when applied to large groups such as broiler (meat) chickens. We here address this issue by showing that camera technology can both provide life-long continuous monitoring of the welfare of whole flocks and also give crucial information about the individuals making up the flock. The cameras detect variation between individuals and are also sensitive to birds moving abnormally. By testing birds individually, we show that slow-moving birds tended to come from flocks that moved slowly overall and showed large variation between individuals whereas fast-moving birds were more likely to come from more active flocks that moved more uniformly. Properly used, camera technology can thus monitor the welfare of flocks continuously throughout their lives while reflecting the behaviour of individual birds.

## 1. Introduction

The use of automated methods for assessing animal welfare is a rapidly growing feature of livestock agriculture [1–5], but commercial poultry farming has raised particular problems because of the large numbers of animals involved. The practical problems of identifying, tracking and locating the many thousands of animals found in large commercial poultry flocks has led to the development of automated systems that do not identify animals as individuals and instead give welfare outcomes that apply to whole groups. For example, flock level analyses of visual images [6–10] and flock sounds [11, 12] deliver useful information on the state of the flock as a whole, not on individual animals. However, such group level approaches to welfare assessment have been criticized on the grounds that they overlook the most crucial element of all – the welfare of individual animals [13, 14]. The aim of this paper is to show that even without specifically identifying individuals, group level methods of welfare assessment can still contribute directly to the welfare of individual animals.

Firstly, group level automated measures of welfare assessment are best seen not as substitutes for human care but as an aid that enables a stockperson to have 24/7 information about their animals and to have their attention immediately drawn to problem areas that need more detailed human inspection or intervention. Their use as an extension to the work of a good stockperson therefore has the potential to lead to an increase in the welfare of individual animals even where the automated system itself does not distinguish between individuals.

Secondly, because automated systems have the capacity to collect much more detailed and more continuous information than is possible for a human observer, they can also do more than simply record the average behaviour of an entire group. Optical flow patterns made by broiler chicken flocks, for example, contain information not only about the average or mean amount of movement within a flock but also how much variation there is in that movement [6, 17]. Such variation is an important part of assessing the state of individual animals within a flock and can be described in a range of ways in addition to a simple measure of variance. The skew of a distribution, for example, is a measure of whether the mode is above or below the mean [15]. The skew statistic from optical flow patterns of broiler flock movements can therefore be used to indicate whether the majority of birds in a flock are more or less active than average. An even more informative way of describing variation is the kurtosis, which is a measure of whether there are abnormally long ‘tails’ or outliers to a distribution. The kurtosis of optical flow in broiler chicken flocks indicates whether there is a ‘tail’ of an abnormally large number of very fast (or very slow) movement events and so can be an indication of whether the most active (or most inactive) birds are in the majority or form a tiny minority. In broiler chicken flocks, kurtosis reflects the activity of the most active individuals in a broiler flock [16] and is correlated with key welfare outcomes including gait, mortality, pododermatitis and hockburn [17–20]. While not describing the state of each individual in a flock, optical flow can thus indicate the proportion of birds to which the overall flock measures apply.

Our aim in this paper is to further establish the value of optical flow patterns at group level as a contribution to understanding individual bird welfare. We tested the hypothesis that the group level outputs (mean, variance, skew and kurtosis of optical flow) correlate with the behaviour of individual birds as assessed in tests designed to measure their movement as individuals. Two of the tests involved the time taken by an individual bird to move down a runway, either with or without obstacles. The third test was the time taken by a bird to sit down in a shallow water bath (the latency-to-lie test), previously shown to be associated with lameness and poor gait scores [21–23]. Specifically, we predicted that birds that moved most quickly down the runways and remained standing for the longest time in the water test would come from flocks with the highest mean and the lowest skew and kurtosis.

## 2. Materials and Methods

### Ethical considerations

All animals were already being raised as agricultural livestock in Switzerland under the specifications of the welfare label BTS which dominates the Swiss market (over 90% of Swiss broilers are produced under this label). The houses all had enclosed outside areas called winter gardens that the chickens could access through popholes. No popholes were opened until the birds were at least 22 days old, after which it was a BTS requirement that they must be opened if the outside temperature was at least 13°C (days 22-29) and 8°C (from day 30 onwards). In addition to the winter garden, the BTS also specifies that there must be a minimum of 15 lux of daylight (which can be supplemented with artificial light) and, from the 10^th^day onward, for the birds to have access to elevated platforms, which increased the available surface by 10%. Cameras were installed in the houses when the houses were empty to avoid disturbance to the birds. The work was approved by the Canton of Bern (BE97/16) and met all cantonal and federal regulations for the treatment of animals.

### Animals and farms

We selected 3 out of 5 farms that were used in a previous study [19] and that had a history of both *Campylobacter* positive and negative flocks according to tests at the abattoir. The 3 farms belonged to one company (Bell AG, Switzerland) and produced broilers under a label with enhanced welfare standards (platforms and wintergardens as described above). Chicks (Ross 308) were placed ‘as hatched’ (mixed sex) as day-olds and grown to a maximum stocking density of 30 kg/m^2^ when including the surface of the raised platforms.

In total, 20 flocks were tested, although complete optical flow records were obtained for only 18. Flock sizes ranged from 11 934 to 24 000 birds (mean: 18 533.7, STD: 3923.0, n = 20) and birds were grown to an age of 30 (2 flocks), or 36 (14 flocks with and without thinning), or 37 (4 flocks with thinning) days.

### Behaviour tests for individual welfare assessment

All behaviour tests were carried out ‘blind’, that is, before the optical flow results or the results from the abattoir were known. On the day of the test, birds were between 23 and 28 days of age (mean: 25.5, STD: 1.32). First, a conveniently chosen group of about 20 birds was separated from the flock by a catching frame (114 x 114 x 60 cm) inside the barn with visual contact to the flock. Noticeably lame or sick birds were excluded. Sixteen randomly chosen birds from this catching frame were marked with color on the head, wings, or tail and one after the other underwent the runway tests. After the tests, chicks were weighed, sexed, and scored for pododermatitis and hockburn while the observer was unaware of the speed in the runway tests. Individual faecal samples were collected before the chick was returned to the catching frame. When all broilers had completed the runway tests the water test was performed in four batches of 4 birds each.

### Runway tests

The length of time a chicken takes to move down a runway towards conspecifics has been used both as a measure of social attraction and also of physical ability to move, particularly where chickens have to overcome obstacles [23–25]. The runway used here consisted of a 342 cm long runway with opaque sides except the far end side where the chicks in the catching frame were visible (Fig. 1). The opaque cover at the side ended one bird length before the catching frame and this was taken as the finish line. One chick at a time was carried to the end of the runway and then released. The time to reach the finish line was recorded with a timeout of 5 min. Immediately afterwards, the test was repeated by adding a line of bricks 14 cm high across the width of the runway and again the time to reach the finish line was recorded with a timeout of 5 minutes.

**Figure 1.**
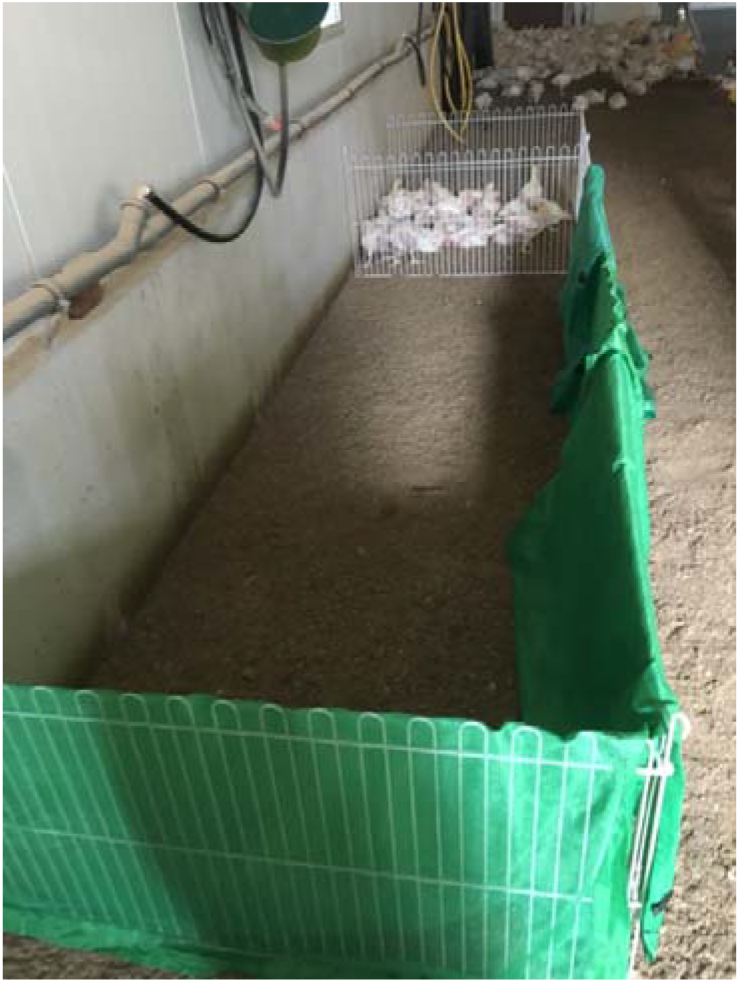
Runway where a chicken was taken from the holding pen (far end) and released at the near end of the runway. The time measured was from release until the chicken came within a bird’s length of its conspecifics in the holding pen, as assessed by the end of the green screen.

### Water test

The method was the ‘latency to lie’ test [21, 22]. Four birds were placed in a box covered with tepid shallow water and watched for 15 min. The time when a bird sat down for the first time was recorded. Broilers that did not sit down were scored with the maximum time of 900 s.

### Optical flow for group level welfare assessment

Two Samsung CCTV IP cameras (SNO-6084RP) were fixed to the ceiling at a height of 5 m on both sides of the barn about one third of the total length of each house away from the entrance. Cameras were installed between flocks to avoid disturbing the birds and were connected to a Synology NAS Disk Station (DS 115j) for video storage via an ethernet switch (HP 9982A). Videos were recorded 24 hours/day from population until a few days before depopulation with 4 images/s, with a resolution of 320 x 240 pixels.

The movements of flocks were analyzed from the output of the cameras by detecting the rate of change of image brightness (‘optical flow’) in different parts of the whole camera image both through time and space [26, 27]. The resulting changes in different parts of an image were then combined to give an estimate of the sum of local velocity vectors. For example, if the entire flock of white chickens on a dark background remained stationary from one frame to the next, there would be no change of brightness and no ‘flow’. But if some chickens moved between frames, some of the white areas would become darker and *vice versa* and this would be registered as a net ‘flow’.

Optical flow can be detected down to pixel level but for reasons of economy, each frame was divided into 1200 (40 x 30) 8-by-8 pixel blocks and the optical flow estimated for each block every 0.25 s. These estimated flow velocities were then combined, on a frame-by-frame basis to give the total ‘flow’ over the entire image expressed as the mean optical flow (indicating overall average movement) plus variance, skew and kurtosis as different descriptors of variation of movement. Further details are given in [20].

To reduce the output to manageable size, the data from 4 frames per second of one camera only were aggregated into sequences of 3600 frames, giving average values of the four optical flow variables (mean, variance, skew and kurtosis) that represented 15 minutes of real time. Median values were used to eliminate spuriously large numbers that occasional occurred in the optical flow records due to artefacts. These 15 minute summaries were then averaged to give the daily (08.00-20.00 hr) values used for the comparison with behaviour tests.

### Statistical analysis

A general linear model (Proc Glimmix, SAS Institute) was used to analyse the runway test without obstacles for the effects of body weight, sex, hockburn and pododermatitis. The time to finish was logarithmically transformed and birds that did not complete the task were deleted. The likelihood of reaching the finish line was analysed using the binary distribution. Residuals were checked for normality. The full model included all interactions apart from those deleted when their P-level was above 0.2.

To compare the behaviour tests with the optical flow results, the behaviour results of the 16 birds selected as a sample from each flock were first combined to give a single average value for that flock for each of the three tests. This resulted in two very different kinds of data – optical flow on whole flocks and behaviour tests on individual birds. To avoid making invalid assumptions about the distribution of data or equality of variances between the two data sets, a non-parametric test of correlation, Spearman Rank Correlation, was used [28].

## 3. Results

### 3.1 Statistical description of flock movement

The optical flow description of the 18 Swiss flocks at 28 days of age is summarized in Table 1. All flocks exhibited a movement distribution with a strongly positive kurtosis, indicating a higher central peak and /or longer ‘tails’ than would be expected in a normal distribution. Kurtosis is measured in standard deviations away from the mean and anything above (or below) 3 constitutes a departure from normality [15]. All flocks also exhibited a positive skew showing that the mode was displaced to the left of the mean, with the ‘tail’ of the distribution to the right.

**Table 1.** Day 28 mean daily optical flow values for 18 flocks with the standard errors in brackets.

| Mean | Variance | Skew | Kurtosis |
| --- | --- | --- | --- |
| 0.24 ($\pm 0.24$) | 0.216 ($\pm 0.215$) | 4.762 ($\pm 4.77$) | 30.109 ($\pm 3.25$) |

### 3.2 Rummy tests

In 7 out of the 20 flocks all broilers reached the finish line within 5 min. in the runway test without obstacles. In the other flocks 1 to 10 out of 16 birds failed to do so. The likelihood to finish the runway test with obstacles decreased with increasing body mass (F_1,282_= 3.93, P = 0.049) and female chicks tended to be less likely to finish the runway with obstacles (F_1,282_ = 3.47, P = 0.064). The same pattern applied to the runway without obstacles but these were non-signficant trends (body mass: F_1,282_= 3.63, P = 0.058; sex: F_1,282_ = 3.42, P = 0.066).

Body mass but not sex, pododermatitis, or hockburn significantly affected the speed to complete the runway test (mass: F_1,236_ = 5.93, P = 0.016; sex: F_1,236_ = 3.06, P = 0.082, pododermatitis: F_1,236_= 0.68, P = 0.41, hockburn: F_1,236_= 2.79, P = 0.10).

### 3.3 Latency to lie (Water test)

Heavier birds of both sexes were more likely to sit down in the water than lighter birds, whereas sex, pododermatitis, and hockburn were not associated with the likelihood (body mass: F_1,275_= 3.97, P = 0.047; sex: F_1,275_= 0.11, P = 0.74; pododermatitis: F_1,275_= 2.27, P = 0.13; hockburn: F_1,275_= 0.08, P = 0.78).

### 3.4 Correlations between individual tests and flock optical flow statistics

The correlations between the three individual behaviour tests and the day 28 optical flow values of the flocks from which the individual birds had been taken are shown in Table 2. In the two runway tests, there was a significant positive correlation between time taken to move through the runway and both the skew and kurtosis of the flocks from which those individuals were taken. The birds that took the most time in the runways came from the flocks with the highest positive skew and the highest kurtosis. In the water test, there was a significant positive correlation between length of time standing and the mean and variance of optical flow. The birds that stayed standing the longest in the water test came from flocks with the highest mean rate of movement.

**Table 2.** Spearman correlations between behaviour tests and optical flow (OF)

|  | Mean OF | Variance OF | Skew OF | Kurtosis OF |
| --- | --- | --- | --- | --- |
| Runway test 1 (The most active birds have the lowest scores) | -0.0781 | 0.008 | **0.711**<br>P < 0.001 | **0.608**<br>P < 0.01 |
| Runway test 2 (with obstacles) | -0.196 | -0.05 | **0.697**<br>P < 0.0025 | **0.625**<br>P < 0.005 |
| Water test (The birds that sit most quickly have the lowest scores) | **0.573**<br>P < 0.01 | **0.574**<br>P < 0.01 | -0.308 | -0.343 |
| Hockburn | -0.021 | 0.147 | **0.508**<br>P < 0.025 | **0.508**<br>P < 0.025 |
| Pododermatitis | 0.023 | -0.003 | 0.398 | **0.426**<br>P < 0.05 |

In the runway tests, 15.5% (Test 1) and 28.5% (Test 2) of the birds did not complete the runway by the end of the test and were scored simply as taking more than 300 s.

In the Water test, 65.5% of birds remained standing at the end of the test and so were scored as taking more than 900 s. The optical flow patterns produced by the movement of broiler chicken flocks showed a positive skew (Table 1), indicating that the mode of the flock movement distribution was displaced to the left and was lower than the mean. All flocks in this study also showed a highly positive kurtosis or right hand ‘tail’ to the distribution, showing that at any one time, there was a small amount of movement that was much higher than the mean activity. Optical flow patterns therefore indicate that broiler chicken flocks consist of a majority of birds that are relatively inactive for most of the time, with a small number of very active birds. This corresponds well with results of studies involving direct behavioural observations that show individual broiler chickens may spend up to 90% of their time sitting or lying and that flocks are typically made up of a majority of inactive birds, with only the minority actively walking or running at any one time [29–31].

## 4. Discussion

The optical flow patterns produced by the movement of broiler chicken flocks showed a positive skew (Table 1), indicating that the mode of the flock movement distribution was displaced to the left and was lower than the median. All flocks in this study also showed a highly positive kurtosis or right hand ‘tail’ to the distribution, showing that at any one time, there was a small amount of movement that was much higher than the mean activity. Optical flow patterns therefore indicate that broiler chicken flocks consist of a majority of birds that are relatively inactive for most of the time, with a small number of very active birds. This corresponds well with results of studies involving direct behavioural observations that show individual broiler chickens may spend up to 90% of their time sitting or lying and that flocks are typically made up of a majority of inactive birds, with only the minority actively walking or running at any one time [29–31].

The results of individual behaviour tests reported here provide further evidence that measures of optical flow-describing movement at flock level are directly indicative of activity shown by different proportions of active and inactive individuals. In the runway tests, the birds that took the longest times to complete the runway came from flocks with the highest skew and kurtosis of flock movement (Table 2). Skew and kurtosis of optical flow are positively correlated with negative welfare outcomes such as mortality, lameness and hockburn [6, 15, 18–21] and the runway tests show that high skew/high kurtosis flocks are the most likely to contain slow moving individuals. High kurtosis is associated with slow movement because in a flock of largely slow-moving birds, very active individuals stand out as an unusual minority and their movements appear as a long ‘tail’ or high kurtosis in the movement distribution. The more birds that are active, the more normal active movement becomes and the lower the kurtosis that is recorded.

In the water test, the shortest latencies to sit down were seen in birds that came from flocks with the lowest mean levels of optical flow (Table 2), providing yet another link between individual and flock behaviour but here with a different optical flow statistic, the mean or average amount of movement. All three individual behaviour tests were thus correlated with optical flow measures at flock level but the correlations were highest with different statistical descriptors. The runway tests were significantly correlated with the skew and kurtosis while the water test correlated significantly with the mean optical flow. This difference can be explained by the differential effects of end-points on the test results. In the runway tests, the birds were given 300 s to move down the runway and 15.% (test 1) and 28.5% (test 2) birds failed to do so within this time. These tests discriminated best between the fastest birds (those that completed the runway in under 300 s), but categorized all the slowest birds as simply taking longer than 300 s. As the slowest birds make up the mode of a flock and therefore contribute most to the overall mean rate of movement, the runway tests were not sensitive to the behaviour of the slowest majority. In the water test, on the other hand, the birds were tested for 900 s. 65.5% of them did not sit down within this time and so were scored simply as taking more than 900 s. The water test is best suited to measuring the performance of the slowest birds (the majority that contributed most to the mean level of movement) but not as sensitive to the behaviour of the most active minority of a flock that would be represented by the kurtosis of the flock optical flow distribution. The different statistical descriptors at group level (mean, variance, skew and kurtosis) are thus all correlated with the behaviour of different individual birds but skew and kurtosis are measures of the activity of the most active members of a flock while the mean or average amount of movement is a measure of the activity of the slower majority.

The average level of activity within a flock has been proposed as a general indicator of the health and welfare of a flock [32] and can easily be automatically detected as mean level of optical flow [6, 7, 10]. However, on its own, average flock activity can be difficult to interpret in welfare terms as it can be influenced by other factors such as breed and light levels. Crucially, it also fails to give any information about the welfare of different individuals within the flock [13]. Our results show that skew and kurtosis of optical flow provide additional powerful information about the proportions of birds within a a flock to which the average flock measures apply. A flock that is active (high mean) but where the most active birds show only slightly more movement than the rest of the flock (relatively low skew and kurtosis) is a flock where most individuals are active. A flock with a high skew and kurtosis on the other hand, indicates a flock where the level of movement of the majority of the flock is below average and healthy active movement is shown of only a tiny minority. Properly used, patterns of optical flow made by broiler chicken flocks thus contain much valuable information about the individuals making up that flock.

## Author Contributions

Conceptualization, Sabine Gebhardt-Henrich, Ariane Stratmann and Marian Stamp Dawkins; Data curation, Sabine Gebhardt-Henrich and Marian Stamp Dawkins; Formal analysis, Sabine Gebhardt-Henrich and Marian Stamp Dawkins; Funding acquisition, Sabine Gebhardt-Henrich and Marian Stamp Dawkins; Methodology, Sabine Gebhardt-Henrich, Ariane Stratmann and Marian Stamp Dawkins; Project administration, Marian Stamp Dawkins; Resources, Sabine Gebhardt-Henrich; Validation, Sabine Gebhardt-Henrich; Writing – original draft, Marian Stamp Dawkins; Writing – review & editing, Sabine Gebhardt-Henrich, Ariane Stratmann and Marian Stamp Dawkins.

Conceptualization, S.G.-H., A.S. and M.S.D.; methodology, S.G.-H., A.S. and M.S.D.; validation, S.G.-H. and M.S.D.; formal analysis, S.G.-H. and M.S.D.; resources, S.G.-H. and M.S.D; data curation, S.G.-H. and M.S.D.; writing—original draft preparation, M.S.D.; writing—review and editing, S.G.-H., A.S. and M.S.D.; project administration, M.S.D.; funding acquisition, S.G.-H. and M.S.D. All authors have read and agreed to the published version of the manuscript.

## Institutional Review Board Statement

Tire study was approved by the Institutional Review Board for the approval of animal experiments of the LANAT office of the Canton of Bern (BE97/16) and met all cantonal and federal regulations for the ethical treatment of animals on 30-09-2016. The procedure was declared to be severity level 0.

## Data Availability Statement

Data collected on farms belongs to the farmers concerned and was made available to us only with their permission. An anonymized version of the data will be made available on application to the authors. In this section, please provide details regarding where data supporting reported results can be found, including links to publicly archived datasets analyzed or generated during the study. Please refer to suggested Data Availability Statements in section “MDPI Research Data Policies” at https://www.mdpi.com/ethics. You might choose to exclude this statement if the study did not report any data.

## Acknowledgments

We would like to thank Lawrence Wang for applying the optical flow algorithm to the video data. We would also like to thank Bell AG for their help with facilitating the project and the farmers and farm managers for letting us put cameras in their chicken houses and allowing us to test their birds and use their abattoir data. Last but not least, we thank Abdulsatar Abdel Rahman for his help with the behaviour tests.

